# In-Vitro Fluorescence Microscopy Studies Show Retention of Spike-Protein (SARS-Cov-2) on Cell Membrane in the Presence of Amodiaquin Dihydrochloride Dihydrate Drug

**DOI:** 10.1101/2021.01.05.424956

**Authors:** Partha Pratim Mondal, Subhra Mandal

## Abstract

The ability of S-glycoprotein (S-protein) in SARS-Cov-2 to bind to the host cell receptor protein (angiotensin-converting enzyme 2 (ACE2)) leading to its entry in the cellular system determines its contagious index and global spread. Three available drugs (Riboflavin, Amodiaquin dihydrochloride dihydrate (ADD), and Remidesivir) were investigated to understand the kinetics of S-protein and its entry inside a cellular environment. Optical microscopy and fluorescence-based assays on 293T cells (transfected with ACE2 plasmid) were used as the preamble for assessing the behavior of S-protein in the presence of these drugs for the first 12 hours post-S-protein - ACE2 binding. Preliminary results suggest relatively long retention of S-protein on the cell membrane in the presence of ADD drug. Evident from the %-overlap and colocalization of S-protein with endosome studies, a significant fraction of S-protein entering the cell escape endosomal degradation process, suggesting S-protein takes non-endocytic mediated entry in the presence of ADD. In contrast, in the presence of Riboflavin, S-protein carries out a normal endocytic pathway, comparable to the control (no drug) group. Therefore, the present study indicates ADD potentially affects S-protein’s entry mechanism (endocytic pathway) in addition to its reported target action mechanism. Hence, ADD substantially interferes with S-protein cellular entrance mechanism. This is further strengthened by 24 hrs study. However, detailed studies at the molecular scale are necessary to clarify our understanding of exact intermediate molecular processes. The present study (based on limited data) reveals ADD could be a potential candidate to manage Covid-19 functions through the yet unknown molecular mechanism.

---

The rapidity with which SARS-Cov-2 is transmitted is unprecedented and the rapid mutation rate resulting in genetic changes, particularly in the spike protein [1], requires special attention to the binding affinity and action mechanism of Spike protein (S-protein) with cell receptor [3]. The ongoing COVID-19 pandemic and many such infections are likely to attract such pandemic emergencies in future with close human-to-human contacts. With rise in global temperature, the situation has become favourable for viruses and bacteria to flourish and emerge in new climate zones where it was never known to exist [4] [5]. Covid-19 may just be the beginning of such pandemic emergencies and frequency of such pandemics are more likely to occur in future.

In the midst of pandemic, we are possibly well-placed to understand the rapidity of transmission, and prevention but not eradication. At the core of this crisis lies the basic understanding of many biological aspects that are currently unknown and the true picture is more likely to emerge in near future. However, the virus SARS-Cov-2 has striking similarities with SARS coronavirus [6]. The host-virus interaction has been well understood, primarily the spike (S) glycoprotein of SARS-Cov-2 that binds with angiotensin converting enzyme 2 (ACE2) on host cell-surface with high affinity primarily by its surface S-glycoprotein [7] [8] [9]. The structural basis of receptor recognition by receptor-binding domain (RBD) of the S-protein of SARS-CoV-2 with the ACE2 reported in details by Shang et al. [10]. The study demonstrated the interaction of ACE2 with RBD of S-protein result in conformational rearrangements of S-protein that could be triggered due to receptor binding with the RBD at cell surface or a change in pH within endosomal environment or both [11].

Present report evaluates S-protein fate upon binding with ACE-2 receptor and estimates how re-purposed drugs may affect S-protein action mechanism. The three re-purposed drugs chosen were Riboflavin [12] [23], Amodiaquin Dihydrochloride Dihydrate (ADD) [17], and Remidesivir [13]. Recent study on bulk whole blood have shown that Riboflavin + UV study significantly could reduce the SARS-CoV-2 titre both in plasma and platelet. However, mechanism of action is still unknown. The ADD second test drug, an FDA-approved drug for malarial and Ebola treatment [16] [17], is also in the list of re-purposed drug that could inhibit SARS-Cov-2 though unknown mechanism [18]. The third re-purposed drug tested is Remidesivir, an established antiviral drug could also inhibit SARS-CoV-2 viral titre compared to control [13]. It is one of the highly exploited drugs for the COVID-19 treatment as it has proven to speed up the recovery times of hospitalized COVID-19 patients [14] [15]. Further, in-vitro and in-vivo study report suggested Remdesivir could be suitable for viral prophylaxis [19] reportedly by inhibition of SARS-CoV-2 virus RNA-dependent RNA polymerase [20], however possibility other mechanism has not been explored. Therefore, lack of understanding about the action of these re-purposed drugs may be the leading cause of evaluating exact potential of these drugs. However, such studies are extremely useful in the middle of a pandemic. Understanding individual proteins rather than complete virus may provide keys to understand virus/host cell receptor binding and its kinetics in the presence of external agent (drugs). As a first preliminary study, present report utilizes above-mentioned three drug candidates to understand kinetics of S-protein upon binding with ACE2-receptor.

Recent studies indicate the efficiency of a specific protein complex (Spike-protein) that binds to the cell ACE2 receptor leading to its entry in target cells [3] [27] [28] [29]. To tackle and manage the virus, several potential neutralizing agents have been proposed but the virus has managed to escape by mutation [30] [31] [32] [33] [34] [36]. Peptide derivatives of ACE2 has shown promise as a potent inhibitor [37]. Our aim is to understand the drug action mechanism at the S-protein to ACE2 receptor level inside a cellular system in the presence of these drugs.

In this brief report, we provide preliminary comparative study of Riboflavin, ADD and Remidesivir drugs effect on the kinetics of S-protein. Precisely, we studied effect of these drugs on the S-protein entry pathway. The study is concentrated on an isolated S-protein of SARS-Cov-2 virus and does not involve the entire virus. So, the results described in this report may be considered limiting in nature.

## I. RESULTS

Fluorescence imaging based investigation has emerged as one of the conclusive evidences for investigating diseases and therapy. The investigation of biological process at the cellular level needed high resolution visualization. Thus, the S-protein mediated host cell uptake mechanism studies described in this report was evaluated based on high resolution confocal microscopy.

### A. S-protein binding study with ACE2-transfected cells

It is well studied that the RBD of the SARS-CoV2 S-protein efficiently binds with human ACE2 expressed on the surface of 293T cells [3] [35]. In the same direction, to study the S-protein binding with ACE2 on cell surface, ACE2 transfected 293T cells were treated with GFP tagged S-protein (GFP-S) and the binding was evaluated over time (Fig. 1). The result shows that GFP-S protein specifically binds with the ACE2 transfected 293T cells (non-transfected cells shows only baseline intensity, as seen in the intensity plots in Fig. 1 bottom row). Furthermore, the 4 hrs and 8 hrs incubation with the S-protein (GFP-S) also illustrated S-protein binds only with 293T cells that had high ACE2 receptors expression (Fig. 1A and 1B), whereas low transfected 293T cells don’t show S-protein binding and have only background intensity (see B1, B2 intensity plots in Fig. 1). However, S-protein seems to uniformly disperse throughout the cell after 12 hrs, as evident from overall reduced intensity throughout 293T cells (see, L1, L2, L3 in Fig. 1). Therefore, the study demonstrates S-protein specific binding affinity with ACE2 receptors on cell surface and overtime gets dispersed within the cells. This is in-line with the studies reported recently by Procko’s group [2] [3].

**FIG. 1:**
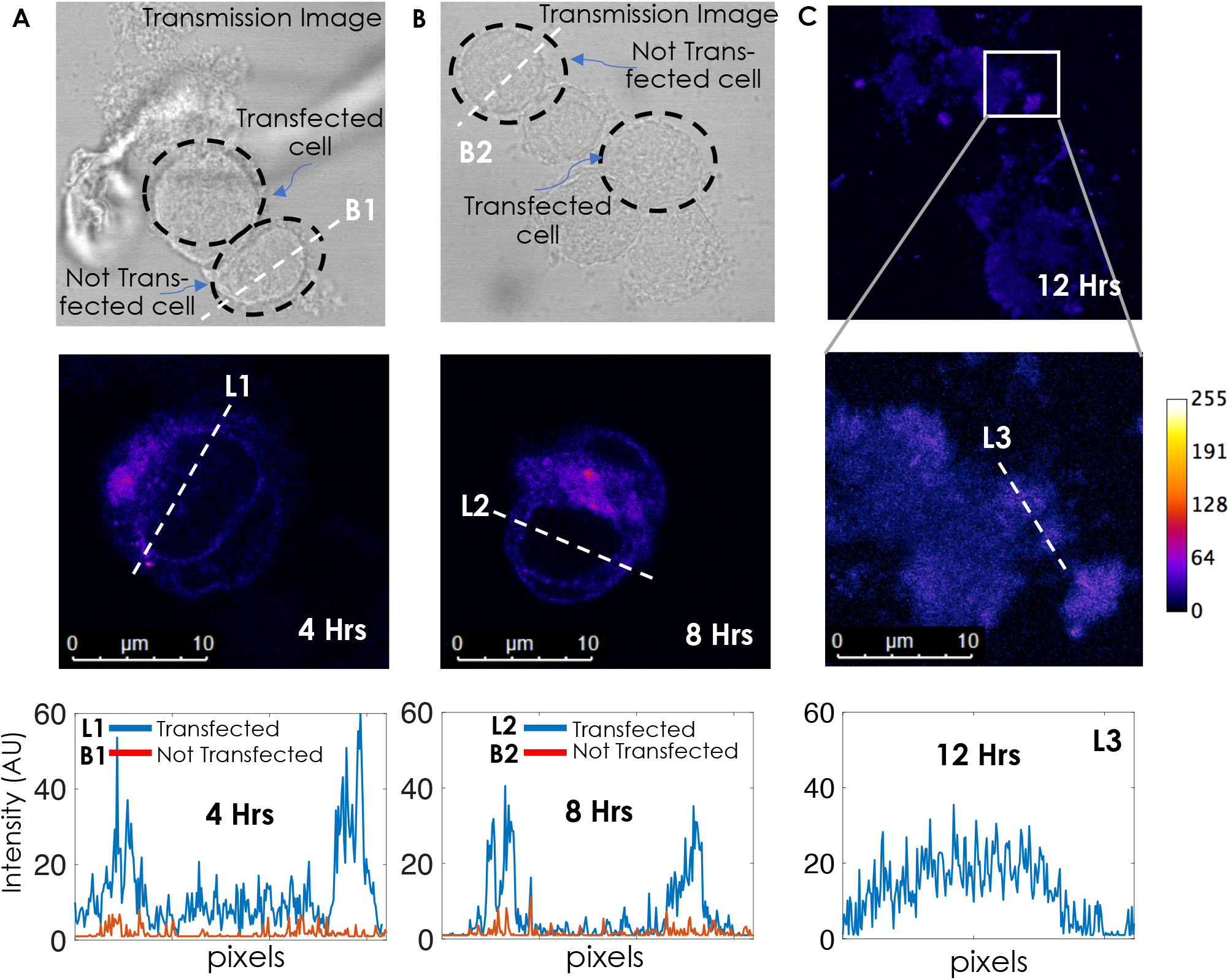
S-protein (GFP-S) binding with ACE2 transfected 293T cells over time. A) 4 hrs and B) 8 hrs GFP-S treated cells. Column represents: Top-Cells under transmission light; Middle-GFP-S/ACE2 positive cell under fluorescent light (excitation: 488 nm; Emission:520 nm); Bottom: Intensity vs pixel plots, L1 and L2 line represent the distribution of S-protein in transfected cell, whereas, B1 and B2 line represent background cell intensity (non-transfected cell). C) 12 hrs GFP-S/ACE2 positive cells under fluorescent light (top), single cell area magnified image (middle), and the intensity vs pixel plots (bottom), where L3 represent the distribution of S-protein in transfected cell.

To comparatively evaluate drug effect on S-protein binding, two sets of experiments were carried out (Fig. 2). In ADD pre-treated set (4 hrs S-protein incubated ADD pre-treated cells, Fig. 2A), the S-protein binding in presence of drug were found predominantly on cellular periphery (Fig. 2A, sky-blue, green and red lines), indicating ADD does not interrupt S-protein binding with ACE2 and S-protein remains adherence on cellular surface. Whereas post ADD-treated cells, i.e., 15 mins ADD treated ACE2 transfected 293T cells incubation with 4 hrs GFP-S followed by 4 hrs GFP-S free fresh medium incubation (8 hrs post ADD treatment, Fig. 2B, sky-blue, green and red lines), indicated S-protein still remained adhered to the cell surface. In both pre- and post-ADD treatment conditions, the S-protein remains bound to the ACE2 transfected 293T cells, demonstrating tested drug (ADD) doesn’t affects the S-protein binding with ACE2 receptors. However, over-time the S-protein shows clustering effect (GFP-S intensity increased, Fig. 2B), indicating possible endosomal accumulation.

**FIG. 2:**
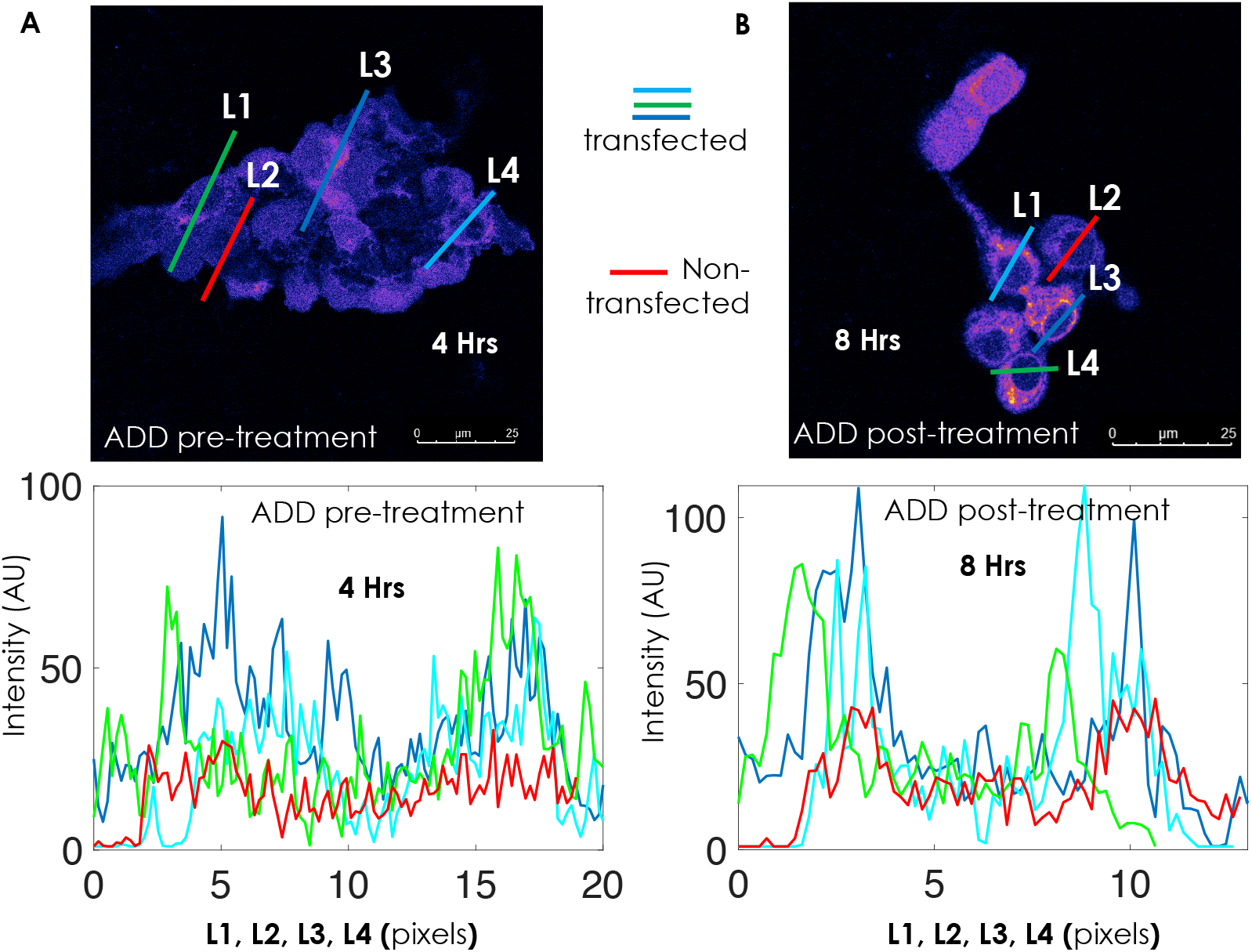
Comparative S-protein (GFP-S) binding study. A) ADD pre-treated (15 mins ADD treatment prior to 4 hrs GFP-S incubation). B) 8 hrs post-treatment: 4 hrs GFP-S incubated ADD pre-treated cells were washed-off and followed 4 hrs in fresh media (with no GFP-S and ADD). Graphs (below) represents fluorescent intensity through the line L1, L3, and L4 (sky-blue, green and red lines) of GFP/ACE2 bound cells vs line L2 (red) for background cell intensity.

### B. Drug effect on S-protein intracellular fate

Next, to evaluate the S-protein (GFP-S) binding and its intracellular pathway mechanism, we followed the GFP-S in early and late endosomal compartments using LAMP-1 antigen/IgG-AlexaFluor 594 lysosome labelled ACE2 transfected 293T cells. Figure 3, demonstrated comparative S-protein (GFP-S, green) binding with ACE2 and its endosomal (LAMP-1, red) fate study of three different re-purposed drug candidates (Riboflavin, ADD and Remidesivir) compared to the control group (untreated). As expected, in case of control (untreated cells), S-protein (green) were endocytosed and enters endocytic pathway (late endosome, red) (Fig. 3, left column) [26]. Similar result was observed in case of Riboflavin and Remidesivir treated cells (Fig. 3, second and fourth column). This demonstrates that the drug treatment doesn’t interferes with the S-protein binding and intracellular uptake mechanism / endocytic pathway (as evident from %-overlap analysis, Fig. 3B). However, ADD shows negligible colocalization illustrating ADD pre-treated condition blocks post S-protein endosomal entrance even after 12 hrs of GFP-S incubation (S-protein not present in endosomes, see Fig. 3, third column). This is evident from the colocalization studies (Fig. 3, fourth row). The study indicates strong proximity of S-protein and endosomes in 293T cells when treated with Riboflavin (Pearson’s correlation coefficient, *r* = 0.51), whereas the least colocalization is reported in the presence of ADD (*r* = 0.19). However, colocalization studies of Remidesivir treated cells (r=0.32) is comparable to untreated control (r=0.33)). In addition, associated intensity plots (L1-L4 in third row Fig. 3 A, and 3B,) also provides a direct evidence supporting above observations. As reflected from 3D cross-section analysis of a cell treated with ADD (Figure 4), it appears that S-proteins are mainly located on the cell membrane post 12 hrs of S-protein - ACE2 binding.

**FIG. 3:**
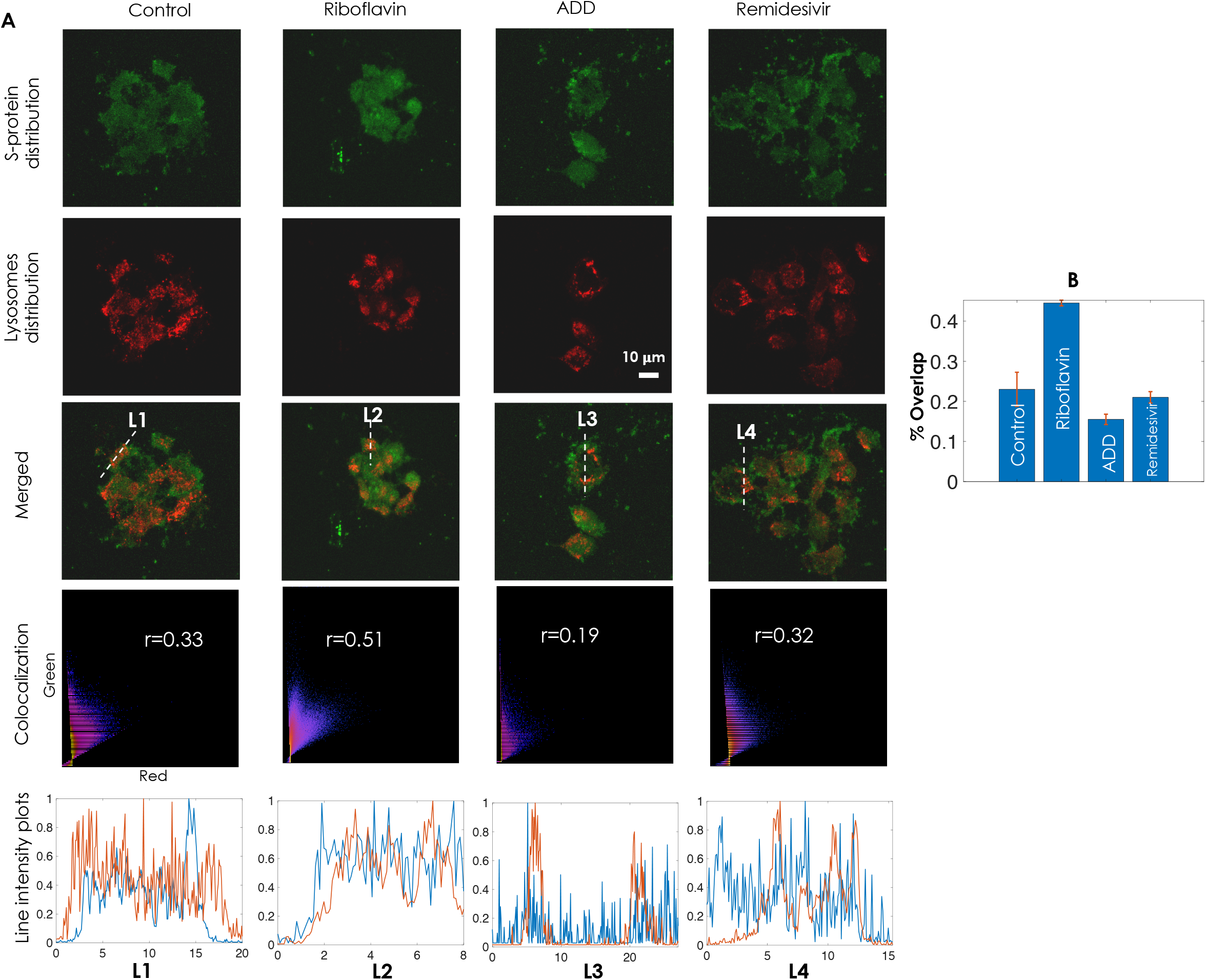
Effect of drugs (Riboflavin, ADD and Remidesivir) on S-protein (GFP-S) distribution and endosomal colocalization study after 12 hours treatment, in comparison to drug untreated control. A) Fluorescent images showing GFP-S (green) distribution and endosomal (red, LAMP1-AlexaFluor 594) colocalization. The colocalization efficiency was evaluated based Pearson’s correlation coefficient, r analysis (4th row) and the line intensity plot (5th row) represent respective line’s fluorescent intensity (GFP-S-blue and Endosome-red) curve. B) The % overlap represent percentage GFP-S protein localization in endosomes in present or absence (control) of treatment.

**FIG. 4:**
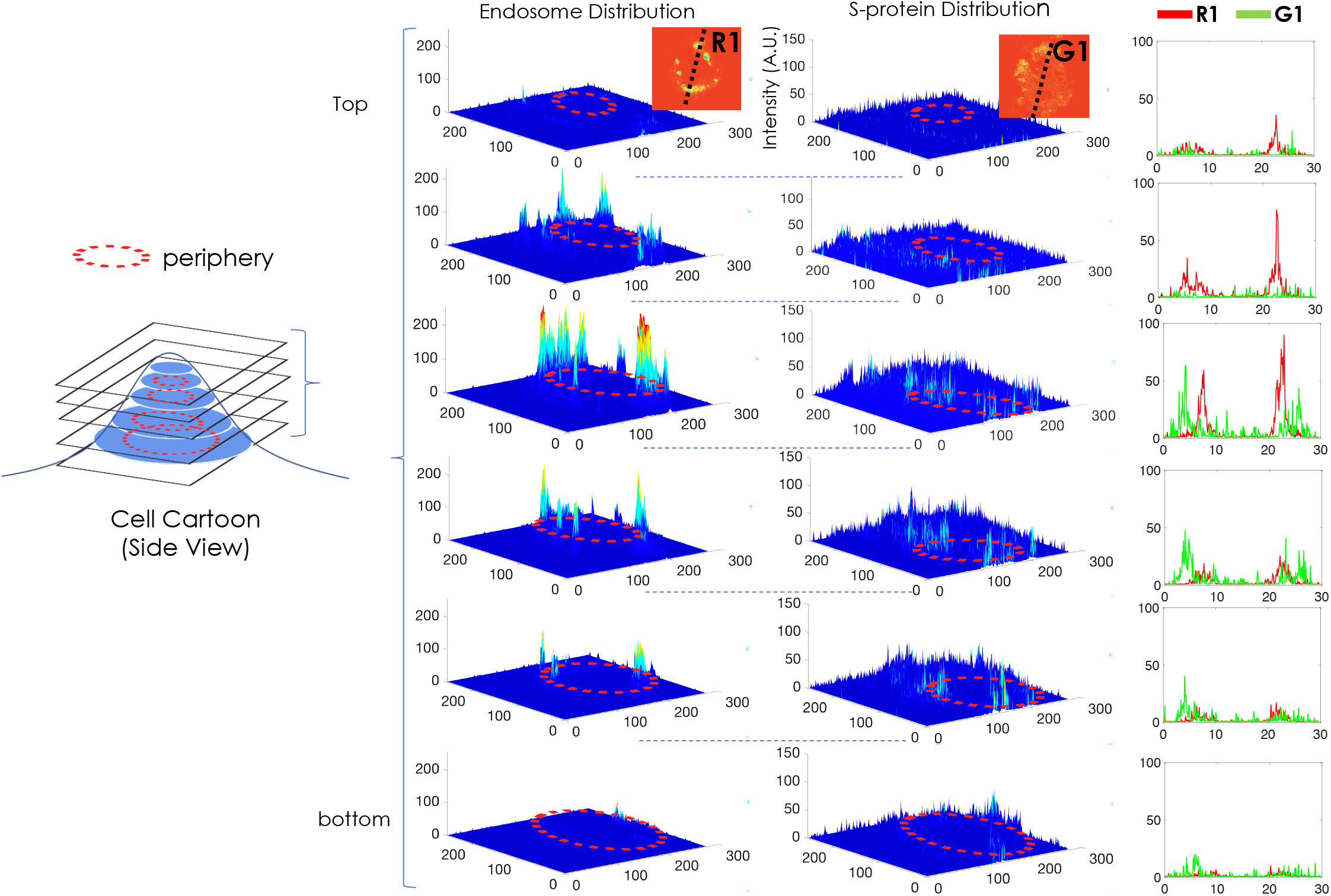
Fluorescent intensity over 3D plane of a 12 hrs ADD treated and GFP-S bound ACE2 transfected 293T cell. A) Side view of a cell cartoon with slices from different planes (top to bottom) is shown. B) 3D surface plots show fluorescent intensity at the respective z-axis surface (all the planes), illustrating the distribution of Endosomes (left column) and S-protein (right column). C) The graphical presentation of fluorescent line intensity of endosome (LAMP1, R1) and S-protein (GFP-S, G1) across the cell in each plane (inset image shows the line across the cell). The line intensity graph shows S-protein (green line) are at the periphery and doesn’t overlap with endosomes (LAMP1, red line).

This is also evident from 24 hrs study (post exposure to drug) as shown in Fig. 5. The corresponding colocalization study (%- overlap analysis in Fig. 5) show a relatively low overlap (between S-protein and Endosomes) for ADD when compared to Remidesivir treatment. Therefore, present study indicated ADD treatment interferes with S-protein cellular endosomal entrance pathway. Additionally, ADD predictively prolongs S-protein retention on the cell surface.

**FIG. 5:**
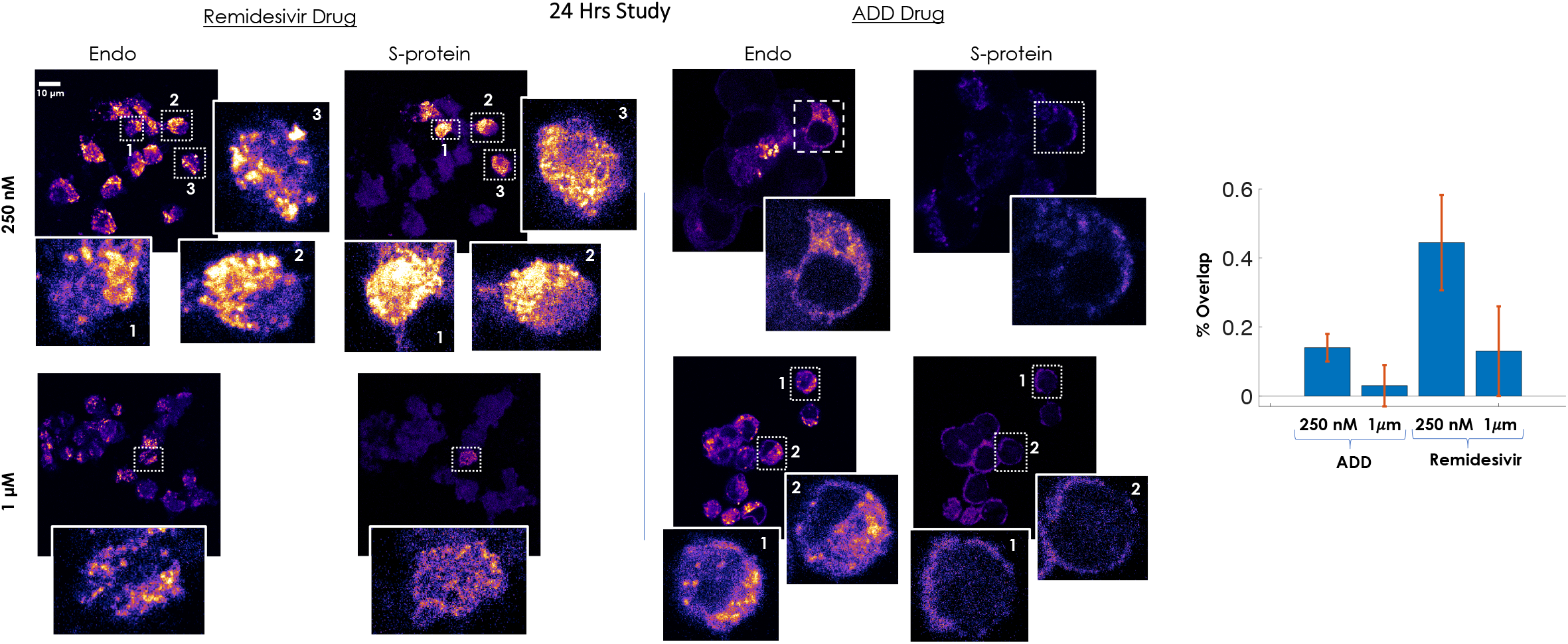
Effect of drugs (ADD and Remidesivir) on S-protein (GFP-S) distribution and endosomal colocalization post 24 hours treatment. Studies are carried out at two different concentrations 250 *nM* and 1 *μM* to access colocalization efficiency. The % overlap represent percentage GFP-S protein localization in endosomes in present of Remidesivir and ADD drugs.

## II. CONCLUSION AND DISCUSSION

The high infection and death rate of SARS-CoV-2 virus has necessitated a quick response to the current pandemic. In the present study, three re-purposed drugs (Riboflavin, Remidesivir and ADD) were investigated to understand their effect on S-protein (SARS-CoV-2 surface protein) kinetics inside and at the cell surface. The ADD, Remidesivir and Riboflavin drugs were chosen due to their reported potency against Covid-19 [17] [23] [19]. Preliminary findings and investigations were carried out that suggest a distinct effect of these drugs on mediated SARS-CoA-2 entry mechanism.

The florescence microscopy-based study indicates ADD drug prolongs S-protein surface retention (Fig. 2) and predictively blocks endosomal entry of S-protein (Figure 3 and 4). The present data reveal that ADD drugs have shown efficacy in Covid-19 treatment, possibly by blocking SARS-CoV-2 endosomal entry (probably preventing pH-dependent S-protein conformational change for needed viral entry [11]), thus prolong retention of the virus on the cell surface. This is further strengthened by long term (24 hrs) study (see, Figure 5). This is important since it gives a longer time for the immune system to act and facilitate medical diagnosis.

In view of the present work being preliminary with limited data, our findings require more investigation and trials for conclusive evidence of the proposed mechanism. We also plan to carry out studies at different time points and with varying concentrations of the proposed drug candidates. In addition, detailed investigation may bring out a clear picture and possible modifications for the mechanisms involved. However, from the present study, it can be speculated that ADD may contribute by moderating the Covid-19 symptoms at this time of pandemic crises.

## III. METHODS AND PROTOCOLS

### A. Study Protocol

In order to investigate the effect of these potential anti-malarial / anti-viral drug candidates on the binding efficiency (of spike protein and cell receptor) and its translation in the cellular system, we developed a simple study protocol as shown in Fig. 6. The central idea is to study both short and moderate time effect of these drugs on 293T cells with S-protein attached to its surface. Step 1 involved the extraction of S-protein primarily based on protocols developed by Procko’s group [3]. The study is broadly categorized into three steps: (1) S-protein extraction, (2) ACE2 receptor transfection of 293 cells, and (3) Drug interaction on S-protein kinetics over time.

**FIG. 6:**
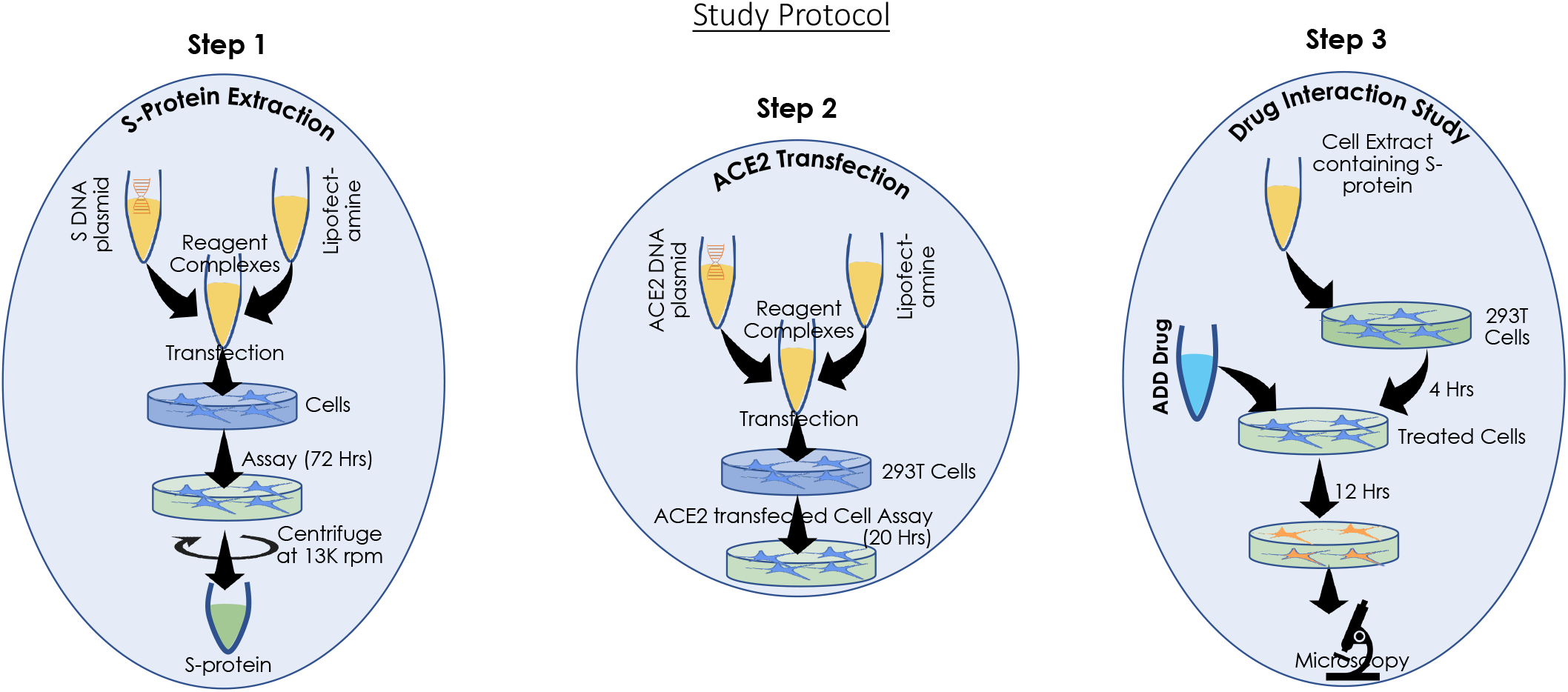
The study protocol for Spike protein extraction, ACE2 transfection and drug interaction study.

The first step towards S-protein study involves the extraction of S-protein through mamallian cell culture. 293T cells were transfected using standard Lipofectamine based protocol. This involves mixing equal proportion of RBD-plasmid (Mammalian expression plasmid for RBD of SARS-CoV-2 protein S (spike) fused with sfGFP) and Lipofectamine to prepare complexes that were added to healthy 293T cells. The protein is secreted. Subsequently, the media from 293T cells secreting the SARS-Cov-2 RBD fused to sfGFP was collected after 72 hrs. The expression medium is then centrifuged to remove cell debris and used for measuring binding. The media containing the protein is directly used as such without purification [2] [3]. No separate protein isolation is carried out.

The next logical step is preparation of 293T cells for drug interaction study. In step 2, a new set of healthy 293T cells were transfected with ACE2 plasmid in a 12-well plate (both with and without coverslip). Coverslip based cell culture and preparation are useful for Confocal microscopy studies. Lipofectamine based protocol with ACE2 plasmid were used to prepare the complexes and added to cells cultured in 12-well plate. In about 20 hrs, about 20 percent of cells were found to be transfected (as seen in fluorescence microscope with blue light excitation) and are ready for next step.

The final step (step 3) is the drug interaction study where these transfected cells were incubated with extracted S-protein medium. The binding efficiency of S-protein (SARS-Cov-2 RBD) labelled with sfGFP was visualized using a blue light (470-490 nm) using a standard inverted fluorescence microscope. Subsequently, the drug is added and these cells (GFP labelled on the cell membrane) were visualized. Finally the cells were fixed at specific time points (12 hrs) and prepared for confocal microscopy studies.

### B. Cell Culture

293T cells were cultured using the standard protocol. Cells were cultured in in a 35 mm and 12-well plate supplemented with medium (DMEM and FBS along with antibiotic (Ampicillin)). The cells were plated at 100, 000 counts and incubated in CO2 incubator. After each 24 hrs, the cell media is removed, cells were washed with PBS to remove dead cells and fresh media was added.

### C. Cell Transfection with ACE2

HEK293T cells were used as host for the present study. Since, HEK293 cells do not express any appreciable level of ACE2 and thus are not susceptible to SARS-CoV-2 infection. Hence, the cells were first transfected with an ACE2 plasmid (name, “pcDNA3.1-hACE2” from AddGene) in order to see binding of the RBD at the surface. Lipofectamine based protocol as described in step 2 of Fig. 5 was used to transfect the cells. For trensfection, antibiotic free medium was used. The cells were incubated for 12 hrs and subsequently the drug interaction study was carried out. This has enabled high efficiency binding of S-protein on host cell surface. These cells are used as a preamble to understand the effect of S-protein-ACE2 binding and its entry in cellular system.

### D. Drug Preparation

The drugs, Amodiaquin dihydrochloride dihydrate and Ribosome were purchased from Sigma Aldritch. DMEM was used to prepare a 250 nM solution of Ribosome, whereas, DMSO is used to prepared 1mM solution of ADD and finally diluted with DMEM to obtain 250 nM solution. In the present study, we have used the above mentioned concentrations although we have seen similar effects at lower concentration as well for which the studies are not reported here. For each study, we have used 200 *μL* of respective drug solution.

### E. S-Protein Extraction using 293T cells as Host

Healthy 293 cells were cultured for extracting S-protein. The cells were thawn and plated in a 35 mm disc with cell medium (95%DMEM + 4.5%FBS + 0.5%Ampicillin). After 4 hrs, the cells were washed with PBS and fresh media was added. The cells were incubated for 48 hrs in a CO2 incubator. A confluency of > 70% is ensured. In the next possage (passage #22) the cells were plated in a fresh 35 mm plate. Following transfection protocol as mentioned in Fig. 1 (step 1), the cells were transfected with SARS-Cov-2 RBD plasmid and left for 72 hrs. The transfected cells were visualization using blue light in a standard inverted fluorescence microscope (Olympus, IX73). The protein is secreted, and the expression medium is centrifuged to remove cells and the medium is used for binding studies. A weak light green color is noted in the cell medium.

### F. S-protein binding and drug treatment method

For the S-protein binding study, the ACE2 transfected HEK293T (293T) cells (Figure 5) were washed and incubated with expression medium containing GFP-S (i.e., GFP expressing S-protein) for 4 and 8 hrs at 37 degrees and 5% CO2. At respective time-point, the cells were washed, fixed by standard protocol (i.e., 30 min incubation in 3.7% formaldehyde at room temperature, treatment wash-off and mounting with FluoroSave, Sigma-Aldrich) and were observed in fluorescence microscope fitted with 20X objective lens and blue light (470-490 nm). For the ADD treatment study, ACE2 transfected 293T were first treated with ADD (250 nM) for 15 mins followed incubated with GFP-S (i.e., GFP expressing S-protein) for 4 and 8 hrs as describe above. At respective time-point, the cells were washed, fixed and were observed in fluorescence microscope as described in pervious paragraph.

### G. Endosome/Lysosome Labelling

For both short and long term (12 hrs and 24 hrs) drug-treatment study, the ACE2 transfected 293T cells were treated with respective drugs, i.e., ADD and Remidesivir (250 nM and 1 *μM*) along with GFP-S containing medium for 12 hrs and 24 hrs in the same condition as mentioned above (see, section F). To visualize Lysosomes / endosomes, LAMP-1 antigen present in late endosomal compartment were targeted. Briefly, the drug treated cells were fixed. The cells were then incubated with the primary mouse anti-LAMP1 antibody (Sigma-Aldrich), at a 50 nM concentration in blocking for 30 min at room temperature. After washing-off primary antibody treatment, the cells were treated with the secondary antibody, Alexa Fluor 594 (emission at 620 *nm*) tagged donkey antimouse IgG antibody (Thermofisher) at 50 nM concentration in blocking buffer for 30 min at room temperature. The cells were then washed 2 times and mounted with FluoroSave before carrying out Confocal studies

## IV. ACKNOWLEDGEMENT

The authors thank Dr. Eric. Procko (University of Illinois, Urbana, IL 61801, USA) for fruitful discussion on hostpathogen interaction. pcDNA3.1-hACE2 was a gift from Fang Li (Addgene plasmid 145033).

## V. DECLARATION

This is to declare that the experimental work was performed at Nanobioimaging Lab, Instrumentation and Applied Physics, Indian Institute of Science, Bangalore, India. We have performed few sets of experiments to report the findings and so further investigations need be performed for conclusive evidence.

